# The closed-loop pathways of signaling molecules

**DOI:** 10.1101/129841

**Authors:** Yang Liu

## Abstract

The pathways of signaling molecules are important to understanding how signaling molecules regulate physiological function and also in predicting the pathological development which is important to therapeutic strategy, however the thorough knowledge of these pathways is still lack. In this paper, we used the big data concept to analyze the pathways of signaling molecules and categorize these molecules into five groups according to their origin and effect on the five organs of heart-spleen-lung-kidney-liver. Heart group includes IGF, Ang and Mg; spleen group includes ANP, aldosterone, retinoic acid and ghrelin; lung group includes FGF7, VEGF, ascorbic acid and HIF; kidney group includes calcitonin, PTHrP, Wnt and NO; and liver group includes EPO, renin, SOD, AKR and GSH. We found that each group of molecules have assisting effect on the other organ in the order of heart-spleen-lung-kidney-liver-heart, and have regulating effect on the other organ in the order of heart-lung-liver-spleen-kidney-heart. Moreover, the pathways of molecules of each group also follow these two arrangements, in which the pathways of molecules form a closed-loop that may lead to new therapeutic strategies.

## 1. Introduction

The pathways of signaling molecules are important to understanding how signaling molecules regulate physiological function and also in predicting the pathological development which is important to therapeutic strategy. However, despite the documented importance of signaling molecules to human life, we still lack thorough knowledge of pathways of signaling molecules. It is well known that heart, spleen, lung, kidney and liver generate/secret important signaling molecules which have significant effect on human physiological processes through signaling pathways. Where liver generates insulin-line growth factor (IGF), heart produces atrial natriuretic peptide (ANP), spleen generates fibroblast growth factor-7 (FGF7) and vascular endothelial growth factor (VEGF), lung secrets calcitonin and parathyroid hormone-related protein (PTHrP), and kidney produces erythropoietin (EPO). These are key signaling molecules, and there are other molecules that regulate or inhibit these key signaling molecules. Here we used Big Data concept to analyze the pathways of these signaling molecules, and categorize these signaling molecules into five groups according to their origin and effect on a particular organ. We named these group as heart group, spleen group, lung group, kidney group and liver group. Heart group includes IGF, Angiotensin (Ang), and Magnesium (Mg); spleen group includes ANP, aldosterone, retinoic acid and ghrelin; lung group includes FGF7, VEGF, ascorbic acid, and hypoxia inducible factor (HIF); kidney group includes calcitonin, PTHrP, Wnt, nitric oxide (NO); and liver group includes EPO, superoxide dismutase (SOD), renin, aldo-keto reductase (AKR), and glutathione (GSH). As shown in Figure 1, within each group, the signaling molecules regulate each other and form a small closed-loop of pathways. Among those five groups, each group have assisting effect on other organ in the order of heart-spleen-lung-kidney-liver-heart, and have regulating effect on the other organ in the order of heart-lung-liver-spleen-kidney-heart. Moreover, the molecules of each group regulate those in other group also in the orders of heart-spleen-lung-kidney-liver-heart and heart-lung-liver-spleen-kidney-heart. Therefore, we may constitute the closed-loop pathways of these signaling molecules.

**Figure 1.**
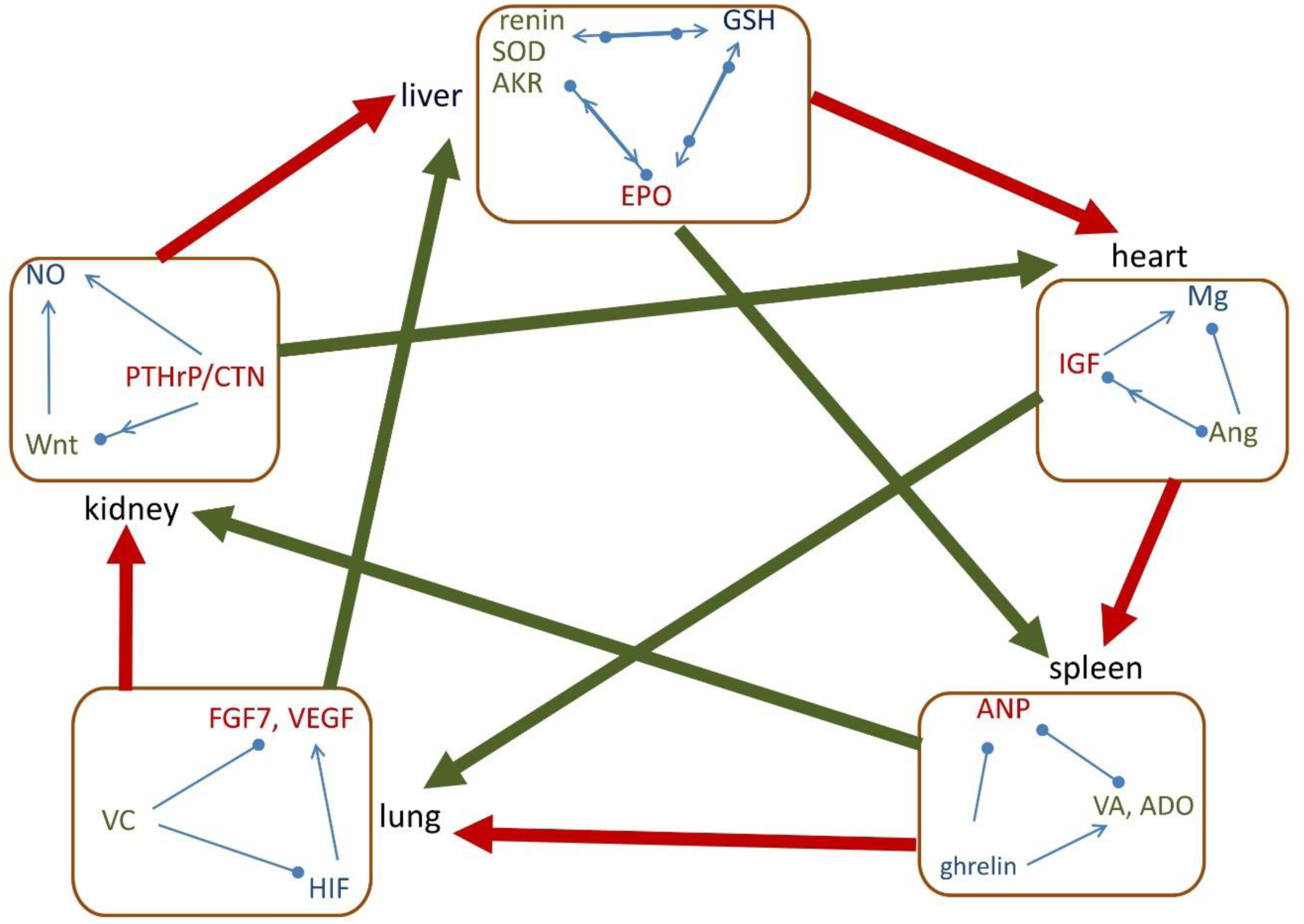
The interrelationships of five group of signaling molecules with organs. The small square indicates the closed-loop pathways of signaling molecules within the same group. Along the outer circle, the thick red arrow shows that the group of molecules assists the following organ, and it follows heart-spleen-lung-kidney-liver-heart arrangement. In the diagonal direction, the thick green arrow shows that the group of molecules assists the following organ, and it follows heart-lung-liver-spleen-kidney-heart arrangement. Arrow means assisting/increasing, solid circle means inhibiting/decreasing, and arrow and circle together indicate regulating.

## 2. Method

We apply the following selection method to identify the signaling molecules.

### Step 1:identify the key molecules

It is known that heart generates ANP, spleen produces FGF7 and VEGF, lung secrets calcitonin and PTHrP, kidney generates EPO, and liver generates IGF respectively. Since IGF has significant effect on heart, ANP has significant effect on spleen, FGF7 and VEGF have significant effect on lung, calcitonin and PTHrP have significant effect on kidney, and EPO has significant effect on liver, we set IGF as key molecule of “heart group”, ANP as key molecule of “spleen group”, FGF7 and VEGF as key molecules of “lung group”, calcitonin and PTHrP as key molecules of “kidney group”, and EPO as key molecule of “liver group”. In Figure 1, along the outer circle of heart-spleen-lung-kidney-liver-heart, the key molecule of each group is generated by the previous organ.

### Step 2: identify the signaling molecules in each group

These key signaling molecules usually have proliferating or warming effect. There should exist other signaling molecules that have shrinkage or cooling effect and also have significant effect on the same organ, i.e., within the same group, the molecules should have contrary effect or regulate each other, thus the functions of these signaling molecules should be

ϕ_1_: proliferation or warming, (for key signaling molecule),

ϕ_2_: shrinkage or cooling,

ϕ_3_: activation or regulation.

Suppose the signaling molecules form the following set,

X = {x: signaling molecules}.

Then, we find the signaling molecules for each group as following set: Heart group= {*x*: ϕ_1_, ϕ_2_, ϕ_3_},

Spleen group = {*x*: ϕ_1_, ϕ_2_, ϕ_3_},

Lung group = {*x*: ϕ_1_, ϕ_2_, ϕ_3_},

Kidney group = {*x*: ϕ_1_, ϕ_2_, ϕ_3_},

Liver group = {*x*: ϕ_1_, ϕ_2_, ϕ_3_}.

Within each group, the molecules form a closed-loop regulating relationship.

### Step 3: Determining the intrinsic relationships among molecules of these five groups

By comparison and iteration, we determine the interrelationships each group of molecules with organs, and the pathways of molecules among these five groups. Thus, we could establish a “large” closed-loop pathways among the molecules of these five groups.

## 3. Results

### 3.1 The signaling molecules in each group and their regulating relationship

The molecules in each group are tabulated in Table 1.

**Table 1.** Groups of signaling molecules

| | $\phi 1$ | $\phi 2$ | $\phi 3$ |
| --- | --- | --- | --- |
| Heart Group | IGF | Ang | Mg |
| Spleen Group | ANP | aldosterone,<br>retinoic acid | ghrelin |
| Lung Group | FGF7, VEGF | ascorbic acid | HIF |
| Kidney Group | calcitonin, PTHrP | Wnt | NO |
| Liver Group | EPO, HGF | renin, SOD, AKR | GSH |

#### 3.1.1 Heart group consists of IGF, Ang and Mg

IGF is secreted by liver and is important to heart function. IGF-1 signaling regulates metabolism, contractility, senescence, autophagy, hypertrophy, and apoptosis in the heart; it activates canonical and noncanonical signaling pathways in the heart; deficiency in IGF-1 may drive cardiovascular disease, and local IGF-1 therapy can prevent heart injuries in experimental models [1]. IGF-1 has warming effect [2].

Ang is secreted by liver; Ang II regulates cardiac and blood vessel contractility, and is involved in cardiac growth, remodeling, and apoptosis [3]. Ang II induces hypothermia [4],which is contrary to IGF.

Mg is important to normal heart rhythm and is a cofactor in more than 300 enzyme systems that regulate diverse biochemical reactions in the body, including blood pressure regulation and blood glucose control [5].

IGF, Ang and Mg regulate each other and form the small closed-loop pathways, as shown in the heart group of Figure 1. It was found that Ang II infusion in rat decreases levels of circulating and skeletal muscle IGF-1 [6], Ang II stimulates cardiac IGF-1 [7] and transcription of IGF-1 receptor in vascular smooth muscle cells [8], and IGF-1 reduces Ang II [9]. Moreover, the exposure of the platelets to Ang II significantly decreases intracellular Mg concentrations [10], and Mg levels are strongly associated with IGF-1 [11].

#### 3.1.2 Spleen group consists of ANP, aldosterone, retinoic acid, and ghrelin

ANP is secreted by heart muscle cells that intervenes in the short-and long-term control of blood pressure and of water and electrolyte balance [12]. The spleen is an important site of ANP-induced fluid extravasation into the systemic lymphatic system. ANP enhances the extravasation of isoncotic fluid from the splenic vasculature both by raising intrasplenic microvascular pressure and by increasing filtration area [13]. Moreover, maintaining body temperature is an important function of ANP [14].

Aldosterone retains needed salt in human body and helps control blood pressure, the balance of electrolytes in blood, and the distribution of fluids in the body. An intracardiac production of aldosterone was recently found in rat [15]. Aldosterone increases spleen size and weight [16]. Retinoic acid (vitamin A) controls the homeostasis of splenic cells [17] and is generated by the epicardium [18]. Moreover, either aldosterone or retinoic acid inhibits heat generation in body [19] [20], which are contrary to ANP.

Ghrelin is a gastric peptide that regulates the distribution and consumption of energy. Ghrelin inhibits proliferation of splenic T cells [21] and effectively regulates food intake and energy homeostasis [22].

The spleen group in Figure 1 shows the contrary effect and supportive relationship between ANP and aldosterone/retinoic acid. ANP has exactly opposite function of aldosterone, ANP decreases circulating aldosterone and vice versa [23]. Retinoic acid signaling markedly stimulates natriuretic peptide receptor-A gene expression [24]. Moreover, it was found that ANP level decreases significantly after treatment with ghrelin in a rat model [25]; ghrelin elevates aldosterone [26]; and applying ghrelin supplementation to normal lungs increased retinoic acid receptor α/γ expression [27].

#### 3.1.3 Lung group consists of FGF7, VEGF, ascorbic acid and HIF

FGF7 is secreted by spleen [28] and is a potent mitogen that enhances cell proliferation in various organs, including the lung, skin, intestine, breast, and liver. It controls the lung morphogenesis, respiratory epithelial cell differentiation and proliferation [29].

VEGF is critical for the development and maintenance of the lung and also plays a role in several acute and chronic lung diseases [30]. It was found that splenic T cells produce VEGF [31]. VEGF is important for brown adipose tissue development and maintenance in which the function of brown adipose tissue is to generate body heat [32].

Ascorbic acid (Vitamin C) was identified in the early 1900s in the search for a deficient substance responsible for scurvy which was directly linked to pneumonia. Vitamin C is a physiological antioxidant protecting host cells against oxidative stress caused by infections, and plays a role on preventing and treating pneumonia [33] and other lung disease [34]. It was found that vitamin C content in spleen is much higher than that in other organs [35] and there exists an extracellular pool of ascorbic acid in lung maintained even during scurvy [36].

HIF which is a highly conserved transcription factor that is present in almost all cell types, is tightly regulated by oxygen availability, and regulates the expression of hundreds of genes. HIF system plays a critical role in pulmonary development [37].

The lung group in Figure 1 shows the contrary and regulating interrelationship of HIF, FGF7, VEGF and vitamin C. HIF-1 activates VEGF transcription in hypoxic cells [38]. HIF promotes the expression of FGF7 mRNA levels [39]. Vitamin C prevents endothelial VEGF and VEGFR-2 overexpression [40] and inhibits NO-induced HIF-1α stabilization and accumulation [41].

#### 3.1.4 Kidney group consists of Calcitonin, PTHrP, Wnt and NO

Both calcitonin and PTHrP are secreted from lung [42] [43]. Calcitonin augments the renal reabsorptive capacity for calcium [44], PTHrP is potent renal regulating factor and mitogenic for various renal cell types, and plays a role in renal development [45]. Both calcitonin and PTHrP have “warming” effect [46, 47].

Wnt signaling regulates cell-to-cell interactions during development and adult tissue homeostasis, and is associated with kidney development and kidney diseases [48]. Several Wnt genes are expressed in lung [49]. Wnt signaling blocks thermogenesis [50] which is contrary to calcitonin and PTHrP.

NO is the second messenger and a major regulator in the cardiovascular, immune, and nervous systems. In the kidney NO has numerous important functions including the regulation of renal hemodynamics, maintenance of medullary perfusion, mediation of pressure-natriuresis, blunting of tubuloglomerular feedback, inhibition of tubular sodium reabsorption, and modulation of renal sympathetic neural activity [51].

The spleen group in Figure 1 shows the interrelationships among calcitonin, PTHrP, Wnt and NO. Calcitonin stimulates expression of sclerostin [52] which is an inhibitor of Wnt, and administration of PTHrP can increase Wnt signaling [53]. Moreover, calcitonin increases plasma NO levels [54], PTHrP activates NO release [55], and Wnt-5a increases NO production [56].

#### 3.1.5 Liver group consists of EPO, renin, SOD, AKR, and GSH

EPO is produced principally in the liver during fetal gestation and mainly in the kidney for adult [57]. EPO is a glycoprotein that promotes the proliferation and differentiation of erythrocyte precursors. It was reported that high-dose EPO increases liver regeneration by affecting the biochemical, morphological, and histopathological parameters after liver resection [58]. EPO increases brown fat gene expression [59].

Renin is produced in kidney and exerts significant effect on liver. The renin-substrate concentration in patients with liver disease is much lower than that of normal subjects [60]. Blockade of the renin-angiotensin system improves liver regeneration [61], i.e., renin is not favor liver proliferation.

SOD is an important antioxidant defense in nearly all living cells exposed to oxygen and is intensively expressed in kidney [62]. SOD mimetic improves the function, growth and survival of small size liver grafts after transplantation in rats [63] and Cu/Zn SOD is broadly distributed in liver [64]. It was found that cold-exposure activates brown adipose tissue thermogenesis but decreases SOD expression [65], indicating SOD has “cooling” effect.

AKR is expressed in kidney [66]. The AKR superfamily comprises of several enzymes that catalyze redox transformations involved in intermediary metabolism, detoxification and biosynthesis [67]. AKR-7A protects liver cells and tissues from acetaminophen-induced oxidative stress and hepatotoxicity [68]. Cold-exposure suppresses AKR level significantly [69], indicating AKR has “cooling” effect.

GSH is a substance produced naturally by the liver and effectively scavenges free radicals and other reactive oxygen species [70]. GSH plays a key role in the liver in detoxification reactions.

The liver group in Figure 1 shows the interrelationships among EPO, renin, SOD, AKR and GSH. When treated with EPO, the GSH and SOD levels in kidney of rats are significantly decreased [71]. The extracellular SOD is a major repressor of hypoxia-induced EPO gene expression [72], but treatment with EPO increases vascular expression of SOD1 [73]. SOD and GSH work together to prevent or repair the damage caused by reactive oxygen species. SODs convert superoxide radical into hydrogen peroxide and molecular oxygen, while the GSH peroxidases (GPx) convert hydrogen peroxide into water, and GSH helps the productivity of GPx [74]. GSH and AKR are also interrelated, since GSH levels are reduced in AKR-deficient strains [75]. Furthermore, renin-angiotensin system stimulates EPO secretion [76], and inhibition of GSH increases plasma renin activity [77].

### 3.2 The assisting relationship between molecule groups and organs

Moreover, as shown in Figure 1, we found that molecules of heart group assist spleen, spleen group assists lung, lung group assists kidney, kidney group assists liver, and liver group assists heart. Along the outer circle it follows the assisting sequence of heart-spleen-lung-kidney-liver-heart. We have also found that the regulating sequence of heart-lung-liver-spleen-kidney-heart, i.e., the diagonal direction. However, the inversed direction may have opposite effect.

#### 3.2.1 The assisting relationship of heart-spleen-lung-kidney-liver-heart

The signaling molecules of heart group have assisting effect on spleen. For example, Mg deficiency causes morphological and immunological alterations in the spleen [78]; it was found that localized IGF-1 secretion enhances erythropoiesis in the spleen of murine embryos [79], lower IGF-I status is associated with higher spleen longitudinal diameter[80], and IGF-I is necessary for survival or transition of myeloid progenitors into the spleen[81]; Ang II has a role in the establishment of an efficient T cell response in the spleen [82].

The signaling molecules of spleen group have assisting effect on lung. For example, ANP attenuates Lipopolysaccharides-induced lung vascular leak [83]; Retinoic acid regulates lung morphogenesis [84]; aldosterone may be used as a strategy to increase lung edema clearance [85]; and ghrelin is produced early by the fetal lung and promotes lung growth [86].

The signaling molecules of lung group have assisting effect on kidney. For example, HIF is involved in the regulation of a multitude of biological processes that are relevant to kidney function under physiological and pathological conditions, including glucose and energy metabolism, angiogenesis, erythropoiesis and iron homeostasis, cell migration, and cell-cell and cell-matrix interactions [87]; FGF-7 is part of the signaling pathway controlling collecting system size and nephron number in the kidney during development [88]; and VEGF is required for growth and proliferation of glomerular and peritubular endothelial cells [89].

The signaling molecules of kidney group have assisting effect on liver. For example, PTHrP activates human hepatic stellate cells[90]; calcitonin intensificate the processes of conjugation of bile acids with amino acids taurine and glycine in hepatocytes and canalicular secretion that result in improvement of solubilization properties of the bile, ability of the bile to hold cholesterol in solute state and prevent the formation of calculi in biliary tracts [91]; short time lower dose of NO administration is good for liver, but long time high dose of NO administration is harmful for liver [92]; Wnt signaling contributes to liver physiology and pathology by regulating various basic cellular events, including differentiation, proliferation, survival, oxidative stress, and morphogenesis [93].

The signaling molecules of liver group have assisting effect on heart. For example, low GSH has high risk of cardiovascular disease [94]; treated with EPO, a significant improvement in cardiac function and symptoms can be found [95]; extracellular SOD protects the heart against oxidative stress and hypertrophy after myocardial infarction [96]; and AKR is activated in the ischemic heart [97].

#### 3.2.2 The regulating relationship of heart-lung-liver-spleen-kidney-heart

The signaling molecules of heart group has regulating effect on lung. For example, IGF-1 induces lung fibroblast activation, and blockade of IGF pathway in murine model of lung fibrosis improved outcome and decreased fibrosis [98]; Ang II could mediate the response to lung injury [99]; and Mg deficiency contributes to pulmonary complications [100].

The signaling molecules of lung group has regulating effect on liver. It was found that FGF7 promotes liver regeneration [101]; VEGF promotes proliferation of hepatocytes through reconstruction of liver sinusoids by proliferation of sinusoidal endothelial cells [102]; vitamin C deficiency promotes fatty liver disease development [103] and vitamin C plus E combination treatment is a safe and effective treatment option in patients with fatty liver disease [104]; and HIF plays a role across a range of hepatic pathophysiology [105].

The signaling molecules of liver group has regulating effect on spleen, for example, EPO is necessary for the development of erythroid colonies in spleen [106].

The signaling molecules of spleen group has regulating effect on kidney, for example, ANP can act directly or indirectly (via inhibition of aldosterone biosynthesis) on the kidney to alter sodium transport and may regulate fluid distribution within the extracellular space [107]; Aldosterone is crucial for sodium conservation in the kidney [108]; retinoic acid is required for kidney development [109]; the kidney degrades ghrelin, and increased total ghrelin levels in chronic kidney disease are primarily due to the decreased degradation of ghrelin in the kidney [110].

The signaling molecules of kidney group also have regulating effect on heart organ. For example, NO regulates cardiac function through both vascular-dependent and -independent effects [111]; congestive heart failure is associated with increased circulating calcitonin gene-related peptide (CGRP), but CGRP attenuates the development of myocardial infarction [112]; PTHrP has a protective effect for heart [113]; Wnt controls heart development but is also modulated during adult heart remodeling [114].

### 3.3 The interrelationships among groups of molecules

Figure 2 shows the pathways of the signaling molecules of the five groups. Along the outer circle heart-spleen-lung-kidney-liver-heart, the signaling molecules activate each other and form a clockwise closed-loop. As between liver group and heart group, EPO administration increases plasma IGF-I levels [115],

**Figure 2.**
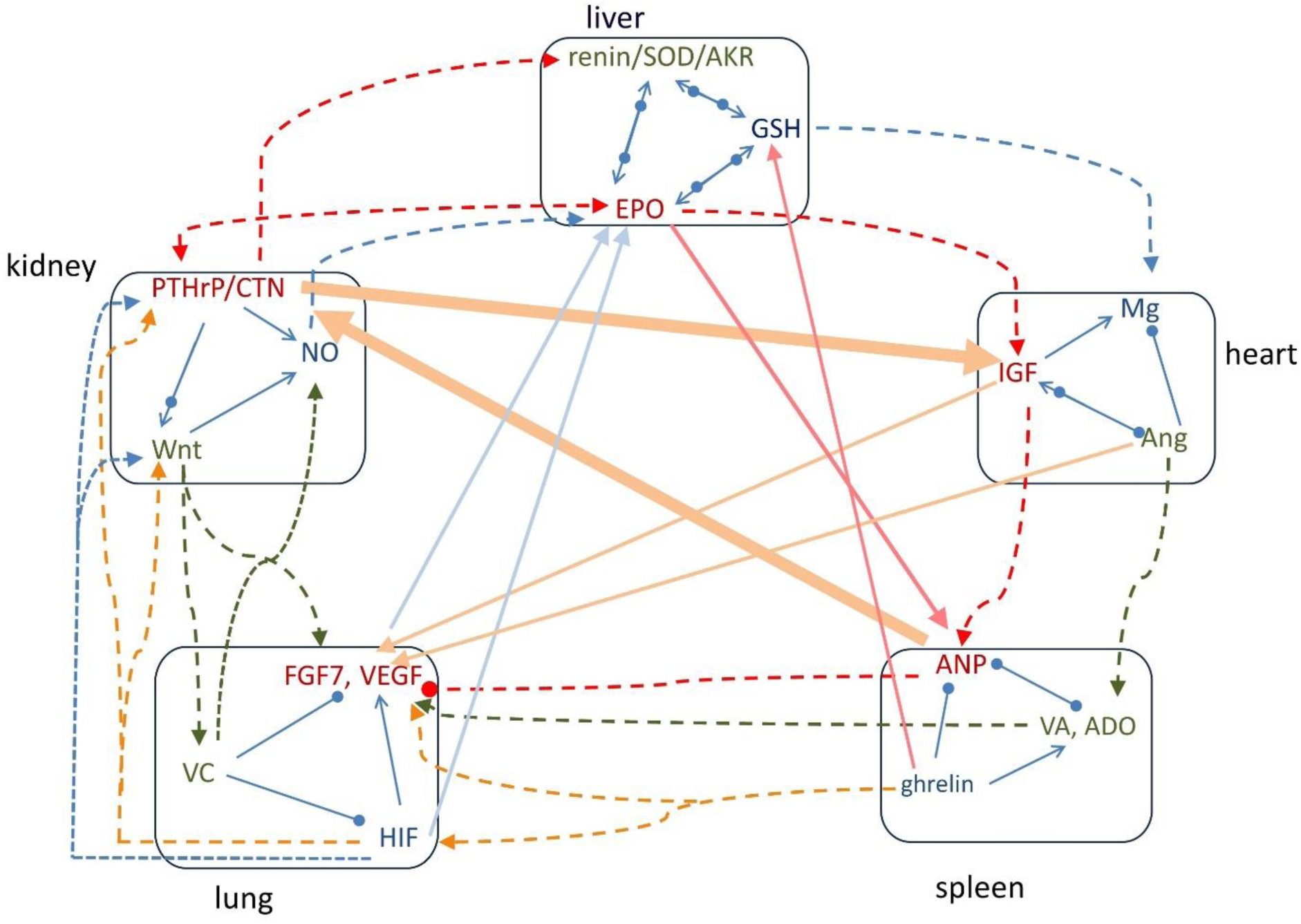
The pathways of signaling molecules among the five groups of molecules. Along the outer circle, the pathways follow the order of heart-spleen-lung-kidney-liver-heart groups. In the diagonal arrangement, the pathways follow the order of heart-lung-liver-spleen-kidney-heart groups. Arrow means increasing, solid circle means decreasing, and arrow and circle together indicate regulating.

GSH enhances intracellular Mg both in vivo and in vitro [116]. Between heart group and spleen group, it was found that IGF-II increases the protein level of ANP [117], and Ang causes the release of aldosterone from the adrenal glands. Between spleen group and lung group, ghrelin administration significantly increased HIF-1α and VEGF mRNAs [118], Aldosterone increases VEGF-A production in human neutrophils [119], and retinoic acid increases HIF-1α [120] and markedly stimulates FGF7 expression [121]; however, ANP blocks VEGF production and signaling in vitro [122]. Between lung group and kidney group, HIF modulates Wnt/β-catenin signaling [123], induces PTHrP [124], and is regulated by NO [125]; Wnt activates FGF7 during epidermal stratification [126], and regulates vitamin C biosynthesis [127]; and vitamin C increases nitric oxide synthase activity [128]. Between kidney group and liver group, calcitonin gene-related peptide (CGRP) and PTHrP stimulate renin secretion [129, 130], PTHrP and EPO are positively correlated with each other[131], PTHrP upregulates AKR 1C3[132], and NO stimulates EPO[133].

For the diagonal arrangement of heart-lung-liver-spleen-kidney-heart, the signaling molecules in each group also enhances those in following group. For example, between heart group and lung group, IGF-1 induces lung fibroblast activation [98] and induces VEGF [134]; and Ang II increases FGF7 mRNA levels [135] and stimulates VEGF synthesis [136]. Between lung group and liver group, VEGF is crucial for EPO-induced improvement of cardiac function [137]; vitamin C elevates red blood cell glutathione in healthy adults [138]; HIF induces EPO production [139] and induction of HIF-1 varies with the GSSG/GSH ratio [140]. Between liver group and spleen group, EPO stimulates ANP secretion [141], and ghrelin treatment increases GSH levels[142] and activity of SOD [143]. Between spleen group and kidney group, ANP increases calcitonin gene-related peptide (CGRP) within human circulation [144]. Between kidney group and heart group, calcitonin increases concentration of IGF [145], and IGF signaling antagonizes the Wnt pathway [146].

## 4. Discussion

Based on the five organs we categorize the signaling molecules into five groups, heart group, spleen group, lung group, kidney group and liver group, respectively. The assisting and regulating relationships follow the outer circle and diagonal arrangement in Figure 1. As shown in Figure 1, along the outer circle arrangement of heart-spleen-lung-kidney-liver-heart, the key molecule of each group is generated/secreted in the previous organ, for example, IGF in heart group is generated in liver, and so on. Within each group, the molecules are contrary but interdependent and regulate each other. Along the outer circle arrangement of heart-spleen-lung-kidney-liver-heart, the molecules of each group have assisting effect on the corresponding organ and the following organ. For example, the molecules in heart group have assisting effect on heart and spleen, the molecules of spleen group have assisting effect on spleen and lung, the molecules in lung group have assisting effect on lung and kidney, the molecules in kidney group have assisting effect on kidney and liver, and the molecules of liver group have assisting effect on liver and heart. If in reversed order, then the key molecule of each group may have negative effect on the previous organ. For example, circulating ANP, the key molecule of spleen group, is greatly increased in congestive heart failure as a result of increased synthesis and release of this hormone [147]; VEGF, the key molecule of lung group, increases mobilization and recruitment of hematopoietic stem cells and circulating endothelial precursor cells to the spleen resulting in splenomegaly [148]; for the key molecules of kidney group, small cell carcinoma of lung is frequently associated with abnormally raised plasma calcitonin [149], and lung cancer is associated with high level of PTHrP [150]; for the key molecule of liver group, patients with polycystic kidney disease had higher levels of EPO [151]. However, IGF of heart group has positive effect on liver, for example the concentration of IGF-1 is low in patients with chronic liver disease mainly due to the decreased liver function [152].

Along the diagonal arrangement heart-lung-liver-spleen-kidney-heart, the molecules of each group also have positive/regulating effect on the following organ. For example, the molecules of heart group have positive effect on lung, the molecules of lung group have positive effect on liver, the molecules of liver group have positive effect on spleen, the molecules of spleen group have positive effect on kidney, and the molecules of kidney group have positive effect on heart. If in reversed order, the key molecule of each may have negative effect on the previous organ. For example, for the key molecule of heart group, higher serum IGF-1 levels were positively associated with chronic kidney disease [153]; for the key molecule of kidney group, calcitonin gene-related peptide suppresses spleen T lymphocyte proliferation [154] and endotoxin induces PTHrP gene expression in splenic stromal cells [155]; for the key molecule of liver group, the correlation between serum-EPO and lung function indices is negative, indicating a negative effect of EPO on lung function [156]. However, the key molecule of spleen group has positive effect on liver, as ANP protects liver from hypoxic injury [157]; and the key molecules of lung group also have positive effect on heart, for example VEGF and its receptors play a role in many different aspects of cardiovascular development, including stem cell differentiation into cardiomyocytes, stem cell migration and survival, and heart development [158], and treatment with VEGF and FGF7 can help to optimize the development of epicardium [159].

The pathways of the signaling molecules also follow the closed-loop arrangement of heart-spleen-lung-kidney-liver-heart. As shown in Figure 2, the molecules in heart group activate those in spleen group, the molecules in spleen group regulate those of lung group, the molecules of lung group regulate those of kidney group, the molecules of kidney group activate those of liver group, and the molecules of liver group regulate those of heart group.

Moreover, particularly the key molecule of each group follows the pathway of heart-lung-liver-spleen-kidney-heart, the diagonal arrangement in Figure 2. The molecules of heart group activate those in lung group; the molecules of lung group activate the molecule in liver group; the key molecule of liver group activate the key molecule of spleen group, but ghrelin in spleen group activates GSH and SOD in liver group; the key molecule of spleen group activates the key molecule of kidney group; and the key molecule in kidney group activates the key molecule of heart group.

In conclusion, we categorized the signaling molecules into five groups according to their origin and effect on organs, and found the intrinsic relationships among these five group of molecules. These intrinsic relationships form the closed-loop of pathways of signaling molecules and follow the particular pathways. This enables us to investigate the effect of balance of signaling molecules on human health and understand the closed-loop of pathways of these molecules, that would lead to new therapeutic strategies and insight into the rules governing physiological processes.

